# The developmental environment mediates adult seminal proteome allocation in male *Drosophila melanogaster*

**DOI:** 10.1101/2025.01.09.632097

**Authors:** Rebecca von Hellfeld, Rebecca Konietzny, Philip D. Charles, Roman Fischer, Benedikt M. Kessler, Stuart Wigby, Irem Sepil, Juliano Morimoto

**Affiliations:** School of Biological Sciences, University of Aberdeen, AB24 2TZ, Aberdeen, United Kingdom; Target Discovery Institute, Centre for Medicines Discovery, Nuffield Department of Medicine, University of Oxford, OX3 7FZ, Oxford, United Kingdom; Department of Evolution, Ecology, and Behaviour, Institute of Infection, Veterinary, and Ecological Sciences, University of Liverpool, L69 7SB, Liverpool, United Kingdom; Department of Biology, University of Oxford, OX1 3PS, Oxford, United Kingdom; Institute of Mathematics, University of Aberdeen, Aberdeen AB24 3UE, United Kingdom; Programa de Pós-graduação em Ecologia e Conservação, Universidade Federal do Paraná, Curitiba 82590-300, Brazil

**Keywords:** Optimality, Condition-dependence, Game theory, sexual selection

## Abstract

Early life conditions can have long-lasting fitness effects on organisms. In insects, crowding during larval stages impose a significant constraint on adult phenotypes due to increased intraspecific competition for resources, which can modulate males’ success in pre- and post-mating competition in adulthood. Evidence for larval crowding effects on male seminal fluid allocation exists but is limited to a small subset of well-known seminal fluid proteins (Sfps), and often overlooks the interactions between male and female phenotypes. We currently lack a comprehensive understanding of how male and female larval crowding interact to affect production, composition, and transfer of the wider seminal proteome. Here, we manipulated *Drosophila melanogaster* larval crowding (low *versus* high) of males and females to generate individuals with large and small body size, respectively. We mated individuals in a fully factorial design, and measured the abundance, composition, and transfer of Sfps. Large males produced Sfps in significantly higher abundances, yet this difference was marginal and not detected when Sfps were analysed individually. Conversely, small males transferred greater quantities of much of their seminal proteome to females than did large males. When analysing proteins individually, 10 Sfps were transferred in significantly higher abundances by small males than large males. Our results suggest that small males invest more on each mating opportunity, potentially as a response of overall fewer mating opportunities due to their reduced size, or due to the larval cues of high population density. This work provides an insight into early life effects on ejaculate allocation in *D. melanogaster* and sheds light on the physiological and behavioural responses to developmental conditions in insects.

## Introduction

Male reproduction was traditionally considered inexpensive compared with female reproduction and thus, not a major constraint on male reproductive success (Bateman, 1948; Dawkins, 2016). This assumption, however, has been overturned by studies showing that sperm and ejaculate resources can be limiting (Dewsbury, 1982; Wedell, Gage and Parker, 2002). Seminal fluid proteins (Sfps) which, together with sperm, are abundant in the ejaculate and play a pivotal role in triggering post-mating responses in the female, can be depleted faster than sperm in the short term, impacting male reproductive success in many species (Hihara, 1981; Sirot *et al*., 2009; Perry, Sirot and Wigby, 2013; Wigby *et al*., 2016, 2020; Sanghvi *et al*., 2024). As a result, it is not surprising that mechanisms regulating Sfp investment have evolved (Cameron, Day and Rowe, 2007; Marcotte, Delisle and McNeil, 2007; Cornwallis and O’Connor, 2009; Reinhardt, Naylor and Siva-Jothy, 2011; Perry, Sirot and Wigby, 2013). The most detailed knowledge of male Sfp allocation comes from *Drosophila melanogaster*, where some studies have shown that males quantitatively and qualitatively modulate the amount of specific Sfps transferred to females (Sirot, Wolfner and Wigby, 2011; Hopkins *et al*., 2019). Sfps in this species have a range of vital roles in reproduction (Swanson *et al*., 2001), for example by governing sperm storage (Neubaum and Wolfner, 1999; Tram and Wolfner, 1999), supporting oogenesis and ovulation (Soller, Bownes and Kubli, 1999; Heifetz *et al*., 2000; Heifetz, Tram and Wolfner, 2001), reducing remating likeliness (Chen *et al*., 1988), and influencing hatching success (Prout and Clark, 2000; Chapman, 2001).

An important ecological factor that can shape males’ ejaculate quality as well as their ability to allocate their ejaculate is early life condition, which has long-lasting fitness implications for individuals (Stevens, Hansell and Monaghan, 2000; McGraw *et al*., 2007; Monaghan, 2008; Morimoto, Pizzari and Wigby, 2016). In insects, crowding during larval development substantially reduces nutrient availability which imposes metabolic and phenotypic constrains to individuals (Than, Ponton and Morimoto, 2020). Individuals experiencing crowded larval environments develop smaller body sizes than individuals from uncrowded larval environments (Monaghan, 2008), which influences, in adulthood, the ability of males to compete for mating opportunities (pre-copulatory sexual selection) and fertilisation (post-copulatory sexual selection) (Amitin and Pitnick, 2007; McGraw *et al*., 2007). Hence, developmental impacts on adult body size may affect how males modulate their ejaculate, considering that larval density affects accessory gland (AG) size, even after accounting for body size (Bretman *et al*., 2016). A study that quantified 2 specific Sfps (sex peptide and ovulin) in *D. melanogaster* found that large males from uncrowded environments, produced these Sfps at higher abundances, but transferred them in lower abundances per mating when compared with small males from crowded larval environments (Wigby *et al*., 2016). This study also found evidence for higher transfer of sex peptide to large (low larval density) females (Wigby *et al*., 2016), highlighting that early life conditions of both males and females can affect ejaculate quality and quantity, and the ability of males to differentially allocate some Sfps in response to their developmental environment (Wigby *et al*., 2016). However, the seminal proteome of *D. melanogaster* is thought to contain at least 292 Sfps (Wigby *et al*., 2020), but to date only two Sfps have been studied with respect to developmental conditions. Hence, while past studies have provided some insights into the dynamics between larval density, body size, sperm, and one or two select Sfps (McGraw *et al*., 2007; Wigby *et al*., 2016; Zeender *et al*., 2023), we currently lack an understanding of proteome-wide changes in seminal fluid due to rearing condition and the social contexts in which matings occur.

To address this problem, we manipulated larval crowding of males and females and used quantitative proteomics to test how early life environment impacts the production, composition and strategic allocation of non-sperm ejaculate components. Proteomics is the ideal technology to better understand how the seminal fluid might be affected by ecological conditions, and it has been a powerful tool to discover novel seminal fluid proteins in *D. melanogaster* (Findlay *et al*., 2008; Sepil *et al*., 2019). We manipulated larval crowding levels to generate distinct body sizes for both males and females: high larval density to produce small adults, and low larval density to produce large adults. We then paired large and small adult males and females in a fully factorial design to compare the abundances of Sfps produced by large and small males and transferred to large and small females. In *D. melanogaster,* this manipulation is known to influence physiology (Kapila, Kashyap, Gulati, *et al*., 2021; Kapila, Kashyap, Poddar, *et al*., 2021; Morimoto, 2022; Morimoto *et al*., 2023), reproduction (Morimoto *et al*., 2017; Narasimhan *et al*., 2023) and the strength of pre- and post-mating sexual selection (Morimoto, Pizzari and Wigby, 2016). Moreover, it also influences male reproductive anatomy (Morimoto *et al*., 2022) and ejaculate traits (McGraw *et al*., 2007; Wigby *et al*., 2016), which has implications for ejaculate composition and sperm competition. More broadly, larval crowding is an ecologically significant trait for *D. melanogaster* (Morimoto and Pietras, 2020) and can provide biological insights into the responses to developmental conditions in other insect species (reviewed by Than, Ponton and Morimoto, 2020). Based on the literature, we predicted that:

1. Sfp production would be influenced by larval crowding, wherein large males would produce higher abundances of Sfps than small males, as was previously observed for select Sfps (Wigby *et al*., 2016); and
2. small males would transfer a higher abundance of Sfps to females, owing to their relative lower mating opportunities compared with large males (Wigby *et al*., 2016). This effect should be particularly strong when small males mate with large females, as these females are perceived to be more attractive for mating (Schlupp, McKnab and Ryan, 2001; Jerry and Brown, 2017). However the potential costs of female attractiveness, arising from increased male courtship harassment and Sfp transfer remain to be determined (Long *et al*., 2009).

## Materials and Methods

### Fly stocks

We used an outbred stock of *D. melanogaster* collected in Dahomey (Benin) in North Africa which has been maintained in large outbred populations (>1,000 individuals) in controlled laboratory conditions with overlapping generations since 1970 (Fowler and Partridge, 1989). All stocks were maintained, and all experiments conducted, at 25 °C on a 12:12 hour (h) light:dark cycle in a non-humidified room. We used a standard sugar-yeast-maize-molasses medium with excess live yeast granules for experiments following a previously published recipe (Morimoto et al., 2017).

### Manipulation of the larval crowding

We followed the protocol established in Morimoto, Pizzari and Wigby, (2016). Briefly, we collected eggs from population cages and manipulated larval crowding levels as following: (a) the high larval density (small bodied adults) had ∼50 larvae/ml of food (∼200 larvae per ∼4 ml fly food in a 34 ml vial) and (b) the low larval density treatment (large bodied adults) had ∼4 larvae/ml of food (∼40 larvae in ∼10 ml fly food in a 34 ml vial). See Morimoto, Pizzari and Wigby, (2016), and Morimoto, McDonald and Wigby, (2023) for full details. These densities also correspond to normal and high larval densities observed in wild populations in *D. melanogaster* (Morimoto and Pietras, 2020). We will herein refer to the body sizes of adult flies as ‘large’ or ‘small’ to indicate individuals reared in low and high larval densities, respectively. Virgin males and females were collected within 6 h of eclosion and kept in in groups of 15-20 individuals within same sex and larval density treatment vials for 5-7 days prior to the onset of experiments.

### Male ejaculate profile

To measure male ejaculate investment in response to the developmental environment, we conducted a fully factorial experiment where large and small males were given the opportunity to mate with large and small females, in all combinations (Figure 1). Mating pairs (*N = 15* per combination) were allowed to interact for 2 h, until mating was observed, after which males that successfully mated were transferred to cryotubes immediately after the end of mating and snap frozen with liquid nitrogen and kept at -80 °C until dissection. Mating pairs (*N* = 3) that failed to mate during the 2 h interaction was excluded. The experiment was replicated independently 5 times (*N total = 300* (*15 mating pairs x 4 mating combinations x 5 replicates*)). We also kept a sample of large and small body size virgin males as non-mating controls (*N = 100* (*10 males x 2 larval treatments x 5 replicates*)). Except for the lack of opportunity to mate, virgin males were treated exactly as the experimental counterparts. We randomly selected 5 males from each treatment (*N* = 30 (*5 males x 6 treatments*)) for dissection. Each sample was kept on ice for 2 minutes (mins), after which males were placed in a glass slide containing phosphate buffered saline (DPBS, pH 7.2, Sigma Aldrich® Cat-No. PC5119). With thin tweezers AG and ejaculatory duct were dissected under a stereoscope. They were then placed in 1.5 ml Eppendorf tubes and frozen at -80 °C for proteomic analysis. AG and ejaculatory ducts of 5 males from the same treatment group and replicate were then pooled for proteomic analysis (*N = 30* (*6 treatments x 5 replicates*)).

**Figure 1.**
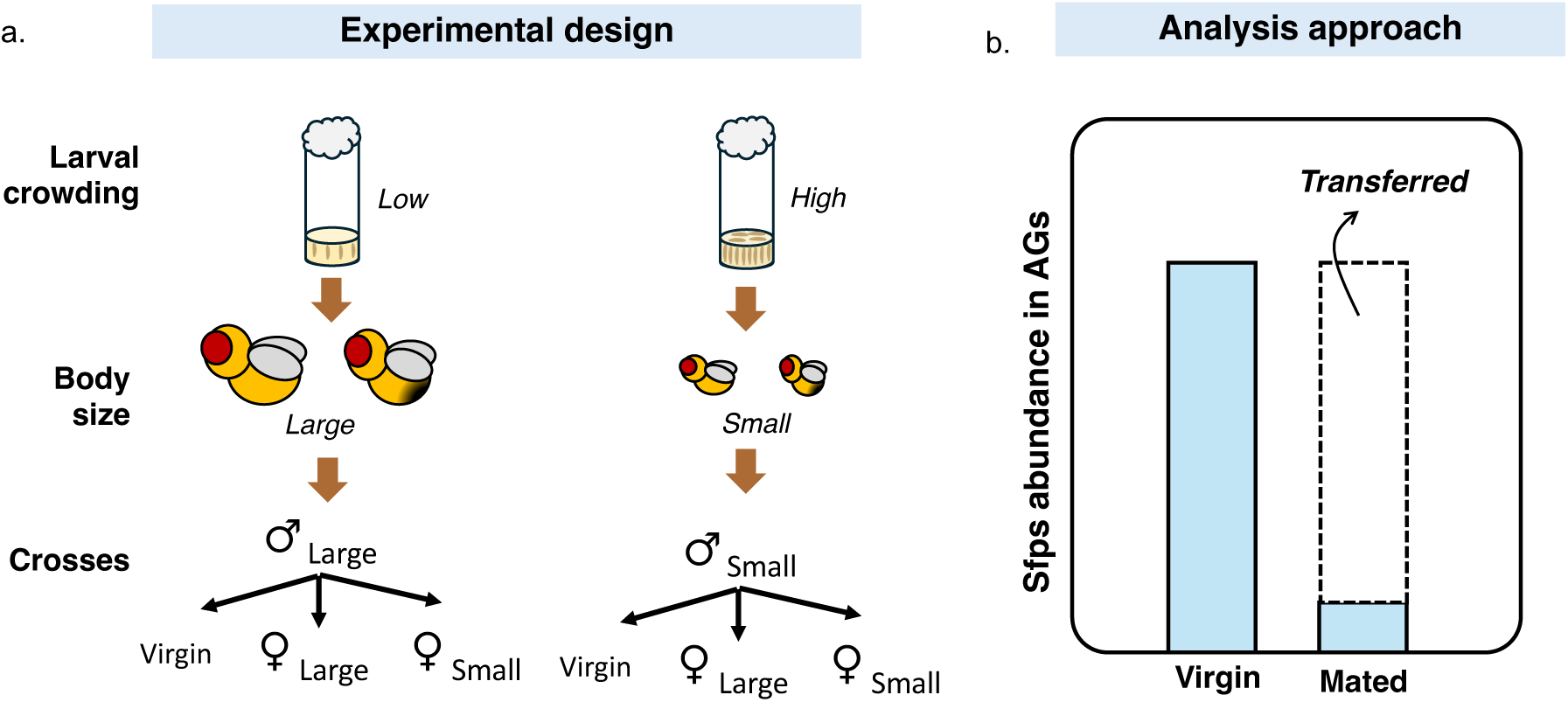
Schematic representation of the experimental design. **(a)** We manipulated larval crowding which resulted in adult flies (both males and females) of large (low crowding) and small (high crowding) body sizes. We then crossed males and females in a fully factorial design. Virgin males of both sizes were maintained to ascertain Sfps production. **(b)** Sfps production was measured as the (standardised) Sfps abundance in AGs of virgin males. Sfps transfer was measured as the difference between the Sfps abundance in virgin males’ AGs minus the Sfps abundance remaining in the AGs of mated males.

### Proteomics sample preparation

Prior to proteomic analysis, we macerated the samples with a clean pestle, washed them with 25 μl Pierce RIPA buffer, and digested the samples using the standard gel-aided sample preparation (GASP) protocol (as described in Hopkins *et al*., 2019; Sepil *et al*., 2019, 2020). First, we reduced the samples by adding 50 mM DTT and waiting for 10-20 mins. Then, we added an equal volume of 40% acrylamide/Bis solution (37.5:1. National Diagnostics) and waited for 30 mins to facilitate cysteine alkylation to propionamide. Next, 5 µl of TEMED and 5 μl of 10% APS were added to trigger acrylamide polymerisation. We shredded the resulting gel plug by centrifugation through a Spin-X filter insert without membrane (CLS9301, Sigma/Corning) and fixed the gel pieces in 40% ethanol/5% acetic acid. This was followed by two rounds of buffer exchange with 1.5 M urea, 0.5 M thiourea, and 50 mM ammonium bicarbonate before removal with acetonitrile. The resulting immobilised proteins were left overnight for trypsin digestion (Promega) and the peptides were extracted with two rounds of acetonitrile replacements the following morning. We dried the peptides using Sola SPE columns (Thermo) and finally resuspended them in 2% ACN, 0.1% FA buffer ready for liquid chromatography-mass spectrometry (LC-MS/MS) analysis.

### LC-MS/MS

The samples were analysed on the LC-MS/MS platform Dionex Ultimate 3000 and Q-Exactive mass spectrometers (Thermo) as previously described in Hopkins *et al*., (2019), and Sepil *et al*., (2019, 2020). First, the peptides were loaded in 0.1% TFA in 2% ACN onto a trap column (PepMAP C18, 300 µm x 5 mm, 5 µm particle, Thermo), followed by peptide separation on an easy spray column (PepMAP C18, 75 µm x 500 mm, 2 µm particle, Thermo) with a gradient 2-35% ACN in 0.1% formic acid and 5% DMSO. MS spectra were obtained in profile mode with a resolution of 70,000 and ion target of 3 x 106. The 15 most intense features were selected for subsequent MS/MS analysis at a resolution of 17,500, a maximum acquisition time of 128 milliseconds, an AGC target of 1 x 105, an isolation width of 1.6Th and a dynamic exclusion of 27 seconds.

### Processing of MS Data

We imported RAW files into Progenesis QIP (version 3.0.6039.34628) using default settings from previous projects and exported MS/MS spectra as MGF files using the 200 most intense peaks without deconvolution for searching (Hopkins *et al*., 2019; Sepil *et al*., 2019, 2020). We chose the *D. melanogaster* UniProt reference proteome as our search target. We used the search engine Mascot 2.5.1. with the following parameters: 10 ppm precursor mass accuracy, 0.05 Da fragment mass accuracy, Oxidation (M), Deamidation (N, Q) and Propionamide (K) as variable modifications, Propionamide (C) as a fixed modification, and two missed cleavage sites. Finally, we applied 1% FDR at peptide level using the target-decoy method inherent to Mascot and an ion score cutoff of 20. The resulting data was imported into Progenesis and the Top3 method was used for protein quantification. The peptide intensity-based quantification data was further cleaned, normalised, and processed as described below.

### Statistical analysis

All analyses were performed using R version 3.0.2 (R Core Team, 2012). First, we removed all the proteins that had less than two unique peptides to avoid incorrect protein assignments. Then, we normalised the data as described by Keilhauer, Hein and Mann, (2015) – a method that we have used previously for normalisation of seminal proteome data (Sepil *et al*., 2019, 2020). The intensity data was initially log transformed [log2(x + 1)] and for each protein a standard deviation of their intensity profile was calculated. Then each protein was ranked according to the standard deviation of their intensity profile and the bottom 90% of the proteins were marked as the ‘background proteome’. This was used to median centre the distribution of each sample.

We ran paired t-tests to compare protein quantities between paired male samples (virgin and mated males from the same treatment and replicate). We applied Benjamini-Hochberg procedure to account for multiple testing. The log2 fold change between the means of the paired male samples and the negative log10 of multiple test corrected p-values were plotted against each other to create volcano plots. Volcano plots were used to visualise the spread of the data and whether the normalisation process was successful. It was also used to check if seminal fluid proteins were significantly depleted following mating, as has been observed in previous projects (Sepil *et al*., 2019, 2020). Proteins were described as Sfps based on the list of 292 published by Wigby *et al*., (2020).

The normalisation process, and the assignment of the background proteome, relies on the assumption that most of the detected proteins have similar intensities between samples. In this dataset for about 90% of the proteins (the background proteome) the standard deviation of their intensity profile was lower than 1.0 [log2(x + 1)] which agrees with this inherent assumption. We also re-ran some of the analyses described below using the unnormalised data to make sure our interpretation of the results is not an artifact of the normalisation process. This additional analysis is reported in Supplementary Information, Figure S1 and S2 and confirms that the results are largely same in both set of analyses.

We focused the rest of our analyses on previously identified Sfps (Wigby *et al*., 2020). The heatmaps were made using a Pearson correlation distance metric and plotted using the ‘pheatmap’ package (Kolde, 2019), and the data were mean-centred (standardised) for each protein for better visualisation.

Major clusters, if found, were determined by visual inspection. The line plots for the clusters were made using the ‘ggplot2’ package (Wickham, 2016), and again used the mean-centred data. Male size-related compositional changes in the seminal fluid proteome and male and female-size related compositional changes in the transferred seminal fluid proteome were assessed using principal component analyses and linear mixed effect models. For Sfp production the initial model included male size as a fixed effect and replicate number as a random effect. For Sfp transfer the initial model included male size, female size and their interaction as fixed effects, and replicate number as a random effect to control for repeated measurements.

Male size-related Sfp abundance differences were analysed for all Sfps together using a linear mixed effect model. Male and female size-related transferred Sfp abundance differences were analysed for each Sfp cluster separately using linear mixed effect models. We inferred the abundance of Sfps transferred to the female by subtracting the Sfp abundance of newly mated males from virgin males within the same treatment and replicate as done previously (Sepil *et al*., 2020). Here, for Sfp production the initial model included male size as a fixed effect, and protein name and replicate number as random effects. For transferred Sfp abundance differences the initial model included male size, female size and their interaction as fixed effects, and protein name and replicate number as random effects. Model selection was performed by backward stepwise elimination. The Database for Visualization and Integrated Discovery (DAVID) was used for gene ontology (GO) enrichment analysis (Huang, Sherman and Lempicki, 2009b).

Male size-related Sfp abundance differences and male and female size-related transferred Sfp abundance differences were also analysed for each Sfp separately using linear mixed effect models. For testing the effect of male size on virgin male Sfp abundances (Sfp production) the initial model included male size as a fixed effect and replicate number as a random effect. For transferred Sfp abundance differences the initial model included male size, female size and their interaction as fixed effects and replicate number as a random effect. Model selection was performed by backward stepwise elimination, and the resulting p-values were corrected for multiple testing using the Benjamini–Hochberg procedure. Sfps that were transferred in significantly different abundances as a response to male size are annotated on the Sfp transfer heatmap, together with Sfps produced in the ejaculatory duct (as opposed to the accessory glands) and Sfps that are known to be functionally important (sex peptide network proteins, ovulin network proteins or proteins linked to sperm competition performance). Finally, we checked whether the functionally important Sfps were over-represented in any of the clusters using proportion tests.

## Results

### Large males produce more Sfps than small males but have similar seminal proteome profiles

From the 30 samples in which five male AG and ejaculatory duct were pooled, we found a total of 2050 proteins. Of these, 1274 had at least two unique peptides and were thus kept in the final dataset. We detected 167 known Sfps out of a total of 292 using the most recent list from Wigby *et al*., (2020). Of the 167 known Sfps, 135 were significantly more abundant in virgin samples than mated samples (for each protein *p* <0.033; 0.166 < log2 fold change < 2.729; Data S1, Figure S3). Only eight Sfps were significantly more abundant in mated *versus* virgin males (for each protein *p* < 0.049; -0.21 < log2 fold change < -0.901). From here on, we focused our analyses on the 167 known Sfps to ascertain if their production and transfer were affected by the size of the male, by the size of its female mate or their interaction.

We first assessed the abundance of Sfps in virgin males in response to our larval crowding manipulation which resulted in distinct body sizes. The heatmap shows no obvious pattern of abundance change within the Sfp proteome in relation to male size and no clear clustering among the Sfps (Figure 2a). However, there is a small but statistically significant change between treatments, whereby Sfp abundances were higher in large *versus* small males overall (F_1,834_ = 8.334, *p* = 0.004; Figure 2b). We investigated whether this effect was driven by some Sfps more than others, by testing for significant changes in individual Sfps. However, we found no evidence for male-size related abundance differences for any of the 167 Sfps in isolation, after multiple test correction (Table S1). These results suggest that large males produce more Sfps than small males, but this effect is not primarily driven by large effects of a small number of Sfp, but instead by relatively small effects in many Sfps.

**Figure 2.**
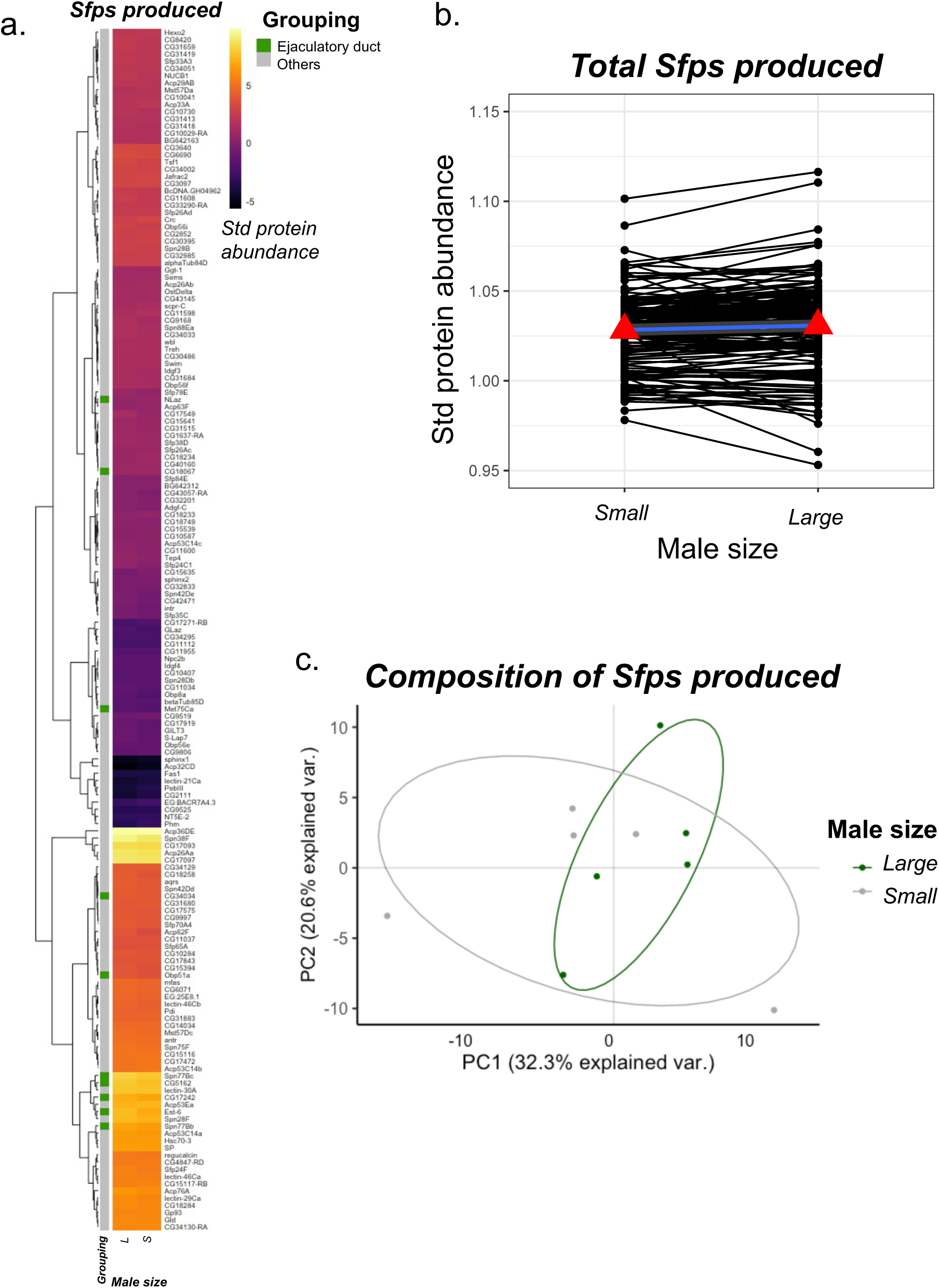
Condition-dependent Sfps production. Seminal fluid protein production increases in large males compared to small males (see also Figure S1 for re-analysis with non-normalised data). **(a)** Heatmap showing the abundance of 167 Sfps identified in accessory gland and ejaculatory duct samples of small and large males. Each cell gives the across-replicate mean for that Sfp in each treatment (n=5 replicates per treatment). None of the Sfps changed in abundance as a response to male size after multiple test correction. Row annotation provide information relating to whether the Sfp is produced in the accessory glands or ejaculatory duct (green annotation). Pearson correlation was used as the distance metric for the hierarchical clustering. **(b)** Line plots showing the change in standardised Sfp abundance with male size. The mean standardised abundance of Sfps in small and large males are depicted with red triangles and joined by a blue line. Large males have a statistically significant increase in Sfp abundances compared to small males. **(c)** Principal component analyses of the seminal fluid proteome in male reproductive tissues. The composition of the seminal fluid proteome is different for small and large males. Green points represent small male virgin samples (5 replicates) and grey points represent large male virgin samples (5 replicates). Ellipses denote 80% normal probability.

Next, we investigated whether the composition of the seminal proteome differed between large and small males, but we did not find any evidence for it (Figure 2c). While larval crowding impacted the abundances of the Sfps, these changes were consistent across Sfps, hence the overall composition remained similar. PC1 explained 32.3% of the variation in the data but this was not associated with the size of the male (L ratio^2^_2_ = 0.768, *p* = 0.381). PC2 explained 20.6% of the variation in the data but again this was not associated with the size of the male (L ratio^2^_2_ = 0.279, *p* = 0.598). We repeated the above analysis with unnormalised data to ascertain whether the patterns were not an artefact of the normalisation process, but the results remained largely unchanged (see Supplementary Information, Figure S1 and S2). Together, these results suggest that although larval crowding strongly influences male size, this translates into minor but widespread differences in Sfp abundances prior to mating.

### Seminal protein transfer responds to male and female developmental environment

Although large and small males have similar Sfp profiles, they may not necessarily transfer similar amounts to females at mating, especially if the females also vary in size. To test this, we investigated how the abundance of transferred Sfps changed between our mating treatment groups. We performed a hierarchical clustering analysis and the resulting heatmap revealed that there was clustering among the Sfps in their response to male and female size. We identified four higher-order clusters and analysed them separately to identify distinct patterns of abundance change within the transferred Sfp proteome (Figure 3a). We also checked whether the distribution of functionally important Sfps among the clusters were biased towards any of them.

**Figure 3.**
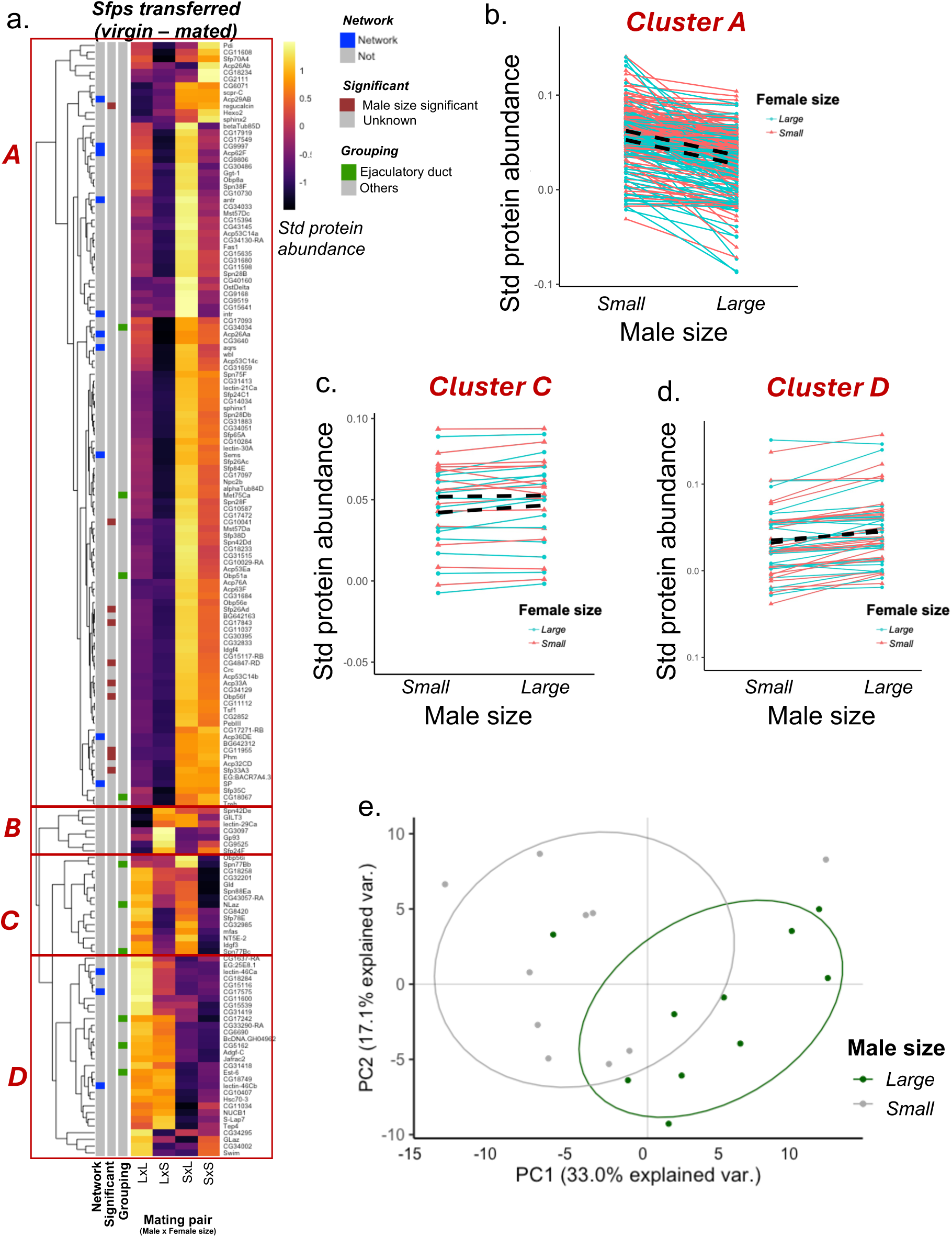
Seminal fluid protein transfer responds to male size when analyses are repeated with unnormalized data. **(a)** Heatmap showing the abundance of 167 Sfps transferred to females during mating (n=5 replicates per treatment). Large males were mated to large females (LxL) or small females (LxS) and small males were mated to large females (SxL) or small females (SxS). Each cell gives the across-replicate mean for that Sfp in each treatment combination. 14 Sfps (Acp33A, CG10041, CG11955, CG17843, CG4847-RD, Obp56f, Phm, Sfp26Ad, Acp53Ea, CG31413, CG34129, PebIII, Sfp26Ac, Adgf-C) were transferred in significantly higher abundances by small males compared to large males. Row annotation provide information relating to whether the Sfp is produced in the accessory glands or ejaculatory duct (green annotation). Pearson correlation was used as the distance metric for the hierarchical clustering. **(b-d)** Line plots showing the standardised abundance of Sfps transferred as a response to male and female size for Sfps in cluster A (b), cluster C (c), and cluster D (d). The average change in Sfp abundance for large and small females as a response to male size are depicted with dashed black lines marked as large F and small F. Small males transferred a higher abundance of these Sfps to large females. **(e)** Principal component analyses of the seminal fluid proteome transferred to females during mating. The composition of the seminal fluid proteome transferred is different for small and large males. Green points represent small male samples (5 replicates mated to small females and 5 replicates mated to large females – 10 in total) and grey points represent large male samples (5 replicates mated to small females and 5 replicates mated to large females – 10 in total). Ellipses denote 80% normal probability.

Cluster A was the largest with 114 Sfps. This group of Sfps responded significantly to male size and female size but there was no interaction between the two (Figure 3b; male size only, L ratio^2^_2_ = 338.15, *p* < 0.0001; female size only, L ratio^2^_2_ = 51.346, *p* < 0.0001; interaction, L ratio^2^_2_ = 0.08, *p* = 0.776). Small males transferred a higher abundance of these Sfps compared to large males. Males also transferred a higher abundance of these Sfps to large females compared to small females. 10 out of 13 functionally important Sfps were found in Cluster A (sex peptide network proteins: antr, intr, CG9997, aqrs, Sems, SP; ovulin network proteins: Acp26Aa; sperm competition performance proteins: Acp62F, Acp36DE, Acp29AB). However, this ratio was not more than expected based on chance (*χ*^2^_1_ = 0.114; *p* = 0.367).

Cluster B was the smallest cluster with eight Sfps (Figure 3a). The Sfps in Cluster B did not respond to male size, female size, or their interaction. The variation in the abundances that were transferred to the female could not be explained by male or female body size (male size: L ratio^2^_2_ = 1.072, *p* = 0.3; female size, L ratio^2^_2_ = 2.404, *p* = 0.121; interaction, L ratio^2^_2_ = 2.204, *p* = 0.137). None of the functionally important Sfps were found in Cluster B, yet this was not different than expected based on chance (*χ*^2^_1_ = 0.011; *p* = 0.543).

Cluster C had 15 Sfps and these were transferred in higher abundances to large females compared to small females (Figure 3c; female size, L ratio^2^_2_ = 16.319, *p* < 0.0001). Male size or the interaction between male and female size did not impact the abundance of the Cluster C Sfps transferred (male size, L ratio^2^_2_ = 1.712, *p* = 0.19; interaction, L ratio^2^_2_ = 1.039, *p* = 0.307). None of the functionally important Sfps were found in Cluster C. Again, this was not different than expected based on chance (*χ*^2^_1_ = 0.369; *p* = 0.728).

Finally, Cluster D was the second largest cluster with 30 Sfps. These Sfps were transferred in higher abundances when the males were large as opposed to small (Figure 3d; male size, L ratio^2^_2_ = 30.848, *p* = 0.0001). The transfer of these was not impacted by female size or the interaction between male and female size (female size, L ratio^2^_2_ = 0.011, *p* = 0.915; interaction, L ratio^2^_2_ = 2.068, *p* = 0.15). 3 out of 13 functionally important Sfps were found in Cluster D (sex peptide network proteins: lectin-46Ca, lectin-46Cb, CG17575). However, this ratio was not more than expected based on chance (*χ*^2^_1_ = 0.007; *p* = 0.465).

For each identified cluster we also conducted a DAVID (Huang, Sherman and Lempicki, 2009b, 2009a) analyses to see if these Sfps were enriched in any function within all the Sfps but did not detect any enriched classes.

The hierarchical clustering analysis and the analyses of the individual clusters revealed that transferred Sfps do not show a uniform response to changes in male and female size. Instead, we found distinct patterns of transfer between protein groups. This suggests that the composition of the seminal fluid proteome transferred during mating might be different depending on male or female size. We tested this with a PCA analysis (Figure 3e). PC1 explained 33% of the variation in the data. For PC1, we observed separate clustering based on male size, indicating that the composition of the seminal fluid proteome transferred is different between large and small males (L ratio^2^_2_ = 8.212, *p* = 0.004). However, the composition of the seminal proteome transferred did not respond to female size or the interaction between male and female size (female size, L ratio^2^_2_ = 2.043, *p* = 0.152; interaction, L ratio^2^_2_ = 0.05, *p* = 0.822). PC2 explained 17.1% of the variation in the data but these were not associated with the size of the male, size of the female or their interaction (male size, L ratio^2^_2_ = 2.114, *p* = 0.149; female size, L ratio^2^_2_ = 0.431, *p* = 0.511; interaction, L ratio^2^_2_ = 0.002, *p* = 0.962). Hence, male size was the primary driver of compositional change in Sfps profiles transferred to females, whereas neither female size nor the interaction between female and male size had an effect.

Finally, we tested whether individual Sfps contributed to this compositional change, to try and identify the key Sfps underpinning compositional effects. After correcting for multiple tests, we found that small males transferred 10 Sfps in significantly higher abundances compared to large males (Table S2). These Sfps are Acp33A, CG10041, CG11955, CG17843, CG4847-RD, Obp56f, Phm, Regucalcin, Sfp26Ad and Sfp33A3 and are all within Cluster A. However, none of the other 135 Sfps were transferred in significantly different abundances based on the interaction between male and female size after multiple test correction (Table S2). Re-analysing the data with raw unnormalized data did not change the results qualitatively (see Supplementary Information, Figure S1 and S2). Together, these results suggest that during mating males have fine control over the quantities of Sfps transferred to females, and small males strategically allocate several Sfps in higher abundance compared to large males and this leads to a compositional change in the seminal fluid proteome transferred.

## Discussion

Our data reveals differential allocation and composition of the seminal proteome based on larval crowding in *D. melanogaster*, driven primarily by males, but also partly by females. We found that large (low larval density) adult males have a small but significant increase in seminal proteome production compared to small (high larval density) males, but that they are compositionally similar. In fact, on an individual basis, no Sfps were observed to be produced significantly more by small males. This finding provides only a weak support for the hypothesis that small males have limited ejaculate resources, compared with large males (prediction 1) (Wigby *et al*., 2016). However, our results broadly support the hypothesis that small males, who are likely weaker pre-mating competitors, can maximise their fitness by investing more Sfps in each mating event, whereas large males, which have more opportunities for mating than small males, could benefit from being more conservative at Sfp allocation to avoid ejaculate depletion (prediction 2) (Wedell, Gage and Parker, 2002; Linklater *et al*., 2007; Sirot *et al*., 2009). However, Sfp allocation at mating was not uniform, suggesting that males invest subsets of Sfps differentially.

We had expected to observe an increase in Sfp production for large males, broadly consistent with previous observations (Wigby *et al*., 2016) and built on the assumption that males with overall more resources would invest more in reproduction (prediction 1) (Dewsbury, 1982; Grafen, 1990; Cotton, Small and Pomiankowski, 2006; De Nardo *et al*., 2021) Our data supported this prediction: large males produced overall a greater abundance of Sfps than small males (Figure 2c). However, the composition of the produced seminal proteome remained unchanged, suggesting that although larval crowding strongly influences male size, this translates into minor but widespread and consistent differences in Sfp abundances prior to mating. This lack of compositional change might suggest that the fitness value of individual Sfps does not change with male size – i.e. Sfps that benefit large males benefit small males in the same way. However, the non-uniform changes to “transferred” Sfps (discussed further below), perhaps argues against this idea. Alternatively, there may be developmental constraints which limit plasticity in seminal proteome composition, especially if large changes in the abundance of individual or groups of Sfps cause a wider reduction in function or effective transfer of the Sfps during mating. This was observed in a previous study where the accumulation of Sfps in the absence of mating led to a change in the seminal proteome composition in old males, yet despite having higher abundances of most Sfps, they were transferred in lower quantities by old unmated males compared to young males (Sepil *et al*., 2020). Moreover, large reductions in a single Sfp, sex peptide, is known to generate widespread changes in the seminal proteome composition and structure, potentially altering many aspects of ejaculate function (Wainwright *et al*., 2021).

Adult male AG size, but not testis size, is affected by both larval crowding, and the presence of adult males during larval development (Bretman *et al*., 2016). AG volumes negatively correlate with increasing larval crowding both in absolute and relative terms, accounting for body size differences (Morimoto *et al*., 2022). AG size has been linked directly to increased production and transfer of sex peptide (Wigby *et al*., 2009), and to increased pre- and post-copulatory success (Bangham, Chapman and Partridge, 2002; Wigby *et al*., 2009). Our proteomics data does not allow us to measure AG volume, but our results suggest that the known differences in AG volume derived from larval crowding likely associate with the abundance of the seminal proteome. Our recent study using microcomputing tomography has shown that AG volume decrease for males that experienced high larval crowding in agreement with sexual selection theory that male AG investment are expected to decrease when the intensity of post-copulatory sexual selection is higher than a given threshold (Morimoto *et. al.,* 2022). However, our past work did not enable us to correlate changes in AG volume with Sfps composition. Future studies combining proteomics with microcomputing tomography will allow us to better ascertain the relationship between AG volume, Sfps yield and compositions directly.

When we assessed the fold-change difference in the abundance of Sfps in the reproductive tract of virgin and mated males, as a measure of Sfp transfer to females, we confirmed prediction 2 that small males transfer higher abundances of Sfps to females. This is in partial agreement with previous literature. Wigby *et al*., (2016) used ELISA to quantify the production and transfer of two Sfps – sex peptide (SP) and ovulin – in a fully-factorial experiment that manipulated larval crowding levels similar to our present design. In their work, they found that large males produced higher abundances of SP and ovulin, but also reported a non-significant trend for large (low crowding) males to transfer proportionally lower abundances of SP in each mating. Supporting this, our proteomic analysis confirmed that large males tend to produce higher abundances of Sfps overall, although we did not have evidence of significantly higher production of SP and ovulin in large *vs* small males. Wigby *et al*., (2016) also reported that males tend to transfer proportionally a higher abundance SP to larger females, although they did not compare the transfer of ovulin across females from different sizes. Our results did not show evidence that large females receive higher abundances of Sfps consistently and the composition of the seminal fluid transferred did not vary as a response to female size. However, Sfps in two of the Clusters A and C had a trend of being transferred in higher abundances to large females. Both SP and ovulin are Sfps from Cluster A, suggesting that there is again partial agreement between our results and the findings of Wigby *et al*., (2016).

One possible explanation for some of the discrepancies is the sensitivity of the methods used to measure Sfps. The ELISA approach by Wigby *et al*., (2016) was highly sensitive to their target Sfps at the expense of global resolution of the seminal proteome. Conversely, our approach was focused on understanding the response of the entire seminal proteome to larval crowding effects. As a result, Wigby *et al*., (2016) approach likely captured relatively smaller changes in the production and transfer of their target Sfps than our method could capture. However, it is important to note that in our study, Sfps transferred were not homogeneous across the seminal fluid proteome, suggesting that males fine-tune and strategically control the transfer of specific Sfps. Specifically, Sfps from Clusters A and C, but not Clusters B and D, were transferred in higher abundances to large females whereas Sfps of Cluster A were transferred in higher abundances by small males compared to large males, but the reverse was true for Sfps in Cluster D. Taken together, the results strengthen the idea that males have control of specific components of their ejaculate and allocate them strategically. Cluster A – the largest cluster – supported the prediction of small males transferring Sfps in a higher abundance compared to large males, and of males transferring a higher abundance to large females (Figure 3). The 10 Sfps transferred in higher abundances by small males compared to large males were: Acp33A, CG10041, CG11955, CG17843, CG4847-RD, Obp56f, Phm, Regucalcin, Sfp26Ad, Sfp33A3. None of these proteins have known or clearly defined reproductive functions. However, Acp33A is associated with sperm competitiveness in *D. melanogaster*: both ‘offence’ (the displacement of previous male’s sperm) and ‘defence’ (protection of stored sperm against displacement) (Fiumera, Dumont and Clark, 2005; Ravi Ram and Wolfner, 2007), and hence might be boosted by small males in anticipation of future or past sperm competition. In particular, the high larval density experienced by small males could be an indicator of high population density, and hence high sperm competition risk (Bretman *et al*., 2016), signalling the need to boost the transfer of sperm competition-related Sfps. The role of Obp56f is unclear, but it does not seem to be required for female fecundity or the induction of female refractoriness after mating (Brown *et al*., 2023). Regucalcin is a Ca^2+^ binding protein which is transferred more by males that mate frequently than by males at their first mating (Sepil *et al*., 2020), however its function as an Sfp also remains unknown.

Once transferred to females, Sfps trigger a wide range of pathways which modulate female immunity, nutrition, vision and overall behaviour (Griffith, 2008; Yapici *et al*., 2008; Häsemeyer *et al*., 2009; Rubinstein and Wolfner, 2013; Sun and Spradling, 2013; Peng *et al*., 2024), through a wide range of communication modes (Kortsmit *et al*., 2024; Zelinger *et al*., 2024). We did not directly measure Sfps in the female reproductive tract and therefore, it will be important for future studies to ascertain whether the reproductive tract of small and large females provide an additional selective barrier that exacerbates or homogenises the response to strategic ejaculate investments from males. This is because we know that larval crowding plays a major role in shaping female sperm receptacle morphology (Amitin and Pitnick, 2007) and that Sfps interacts with and within the sperm receptacle to facilitate sperm storage and release and to trigger a wide range of post-mating behaviours (e.g., Guan, Yu and Li, 2023).

Likewise, it will be important to understand how condition-dependent ejaculate investment, and any putative interactions between Sfps and the female reproductive tract, translate into fitness differences between large and small males mating with large or small females. For *D. melanogaster*, the larval density does not affect male remating intervals, whilst females from low density environments remate more rapidly (Amitin and Pitnick, 2007). A recent study found no differences in mate preference, female mating latency, nor female-male interactions in individuals which experienced diluted larval diets, a treatment which mimics in some ways the developmental effects observed in crowded cultures (e.g. slower growth, delayed development) (De Nardo *et al*., 2021). However, here it was observed that small males transferred more sperm to non-virgin females, displaced a larger volume of previously deposited sperm, and achieved higher paternity share per mating than large males (De Nardo *et al*., 2021). This suggests that smaller males may compensate for reduced mating successes by investing more sperm in each mating event. Our work supports the idea of overall higher ejaculate investment by small males, because we found that small males transferred higher abundances of Sfps to females overall, albeit this trend not being homogeneous across the seminal fluid proteome. Another important point to mention is that our proteomics approach allows to compare relative abundances for a large pool of proteins for which no specific antibodies exist, as is the case for most Sfps. However, our approach is indirect and only provides relative quantification of protein abundances. Future research that builds on our findings by measuring Sfps in absolute quantities, for both production and transfer to females, will provide scope for deeper mechanistic insights.

## Conclusions

Evidence supporting plasticity in sperm and non-sperm production and allocation patterns is ubiquitous (Simmons, 2002; Wedell, Gage and Parker, 2002; Hodgson and Hosken, 2006; Cameron, Day and Rowe, 2007; Birkhead, Hosken and Pitnick, 2009; Wigby *et al*., 2009; Sirot, Wolfner and Wigby, 2011; Perry, Sirot and Wigby, 2013). Using label-free quantitative proteomics, our study shows evidence for condition-dependent ejaculate allocation which supports broad predictions from condition-dependent sexual selection theory in relation to reproductive investment. These findings advance our understanding of how early life ecological conditions can modulate ejaculate allocation in adulthood. It will be important in future to understand the trade-offs associated with the seminal strategies revealed here, and how it relates to sperm allocation, other life history traits, and ultimately fitness. It is likely that larval density effects on male reproductive traits are common amongst insects. Understanding the impacts of larval crowding on ejaculate proteomics could be useful for applications that involve sterile males’ release in biocontrol programmes (Sirot, Wolfner and Wigby, 2011; Wei *et al*., 2016). Optimising developmental conditions to maximise male competitiveness could contribute to the success of such programs. Overall, our findings advance our knowledge of the seminal proteome which is portable to other organisms including humans (Rowe and Houle, 1996; Wigby *et al*., 2020).

