## Supplementary File for "The developmental environment mediates adult seminal proteome allocation in male *Drosophila melanogaster*"

Authors’ affiliations

[a] School of Biological Sciences, University of Aberdeen, AB24 2TZ, Aberdeen, United Kingdom.

[b] Target Discovery Institute, Centre for Medicines Discovery, Nuffield Department of Medicine, University of Oxford, OX3 7FZ, Oxford, United Kingdom.

[c] Department of Evolution, Ecology, and Behaviour, Institute of Infection, Veterinary, and Ecological Sciences, University of Liverpool, L69 7SB, Liverpool, United Kingdom.

[d] Department of Biology, University of Oxford, OX1 3PS, Oxford, United Kingdom.

[e] Institute of Mathematics, University of Aberdeen, Aberdeen AB24 3UE, United Kingdom.

[f] Programa de Pós-graduação em Ecologia e Conservação, Universidade Federal do Paraná, Curitiba 82590-300, Brazil.

Ͳ Present address: Agilent Technologies, Karlsruhe, Baden-Württemberg, Germany.

**Reanalyses of raw data prior normalisation**

One of the main findings in this study is that small males transfer higher abundances of most Sfps in comparison to large males. We wanted to make sure this result was not an artefact of the normalisation process. Due to the differences in male size, small male samples might have a significantly and systematically lower protein content than large males. In such case this true biological variation would be treated as technical variation in the normalisation process and get corrected for by increasing the protein abundances of small males to enable cross sample comparisons. And this could lead one to conclude that small males have and transfer higher abundances than they do - in error.

To make sure this wasn't the case we ran additional analyses and plotted the data using unnormalized abundances. We predicted that if the normalization process was over inflating the protein abundances in small male samples, then the heatmaps of unnormalized data would show inconsistent results where large males produce significantly higher abundances of Sfps and transfer similar or significantly higher abundances of Sfps compared to small males and this would also be evident on the per protein level.

The heatmap of produced Sfps showed no obvious pattern of abundance change within the Sfp proteome in relation to male size and no clear clustering among the Sfps in agreement with the normalised data analyses (Figure S1a). There was a statistically significant change between treatments, whereby Sfp abundances were higher in large *versus* small males overall ($F_{1,834} = 366.3$, $p < 2.2e^{-16}$; Figure S1b). We investigated whether this effect was driven by some Sfps more than others, by testing for significant changes in individual Sfps. However, we found no evidence for male-size related abundance differences for any of the 167 Sfps in isolation after multiple test correction, consistent with the results generated from the normalised data.

Next, we investigated whether the composition of the seminal proteome differed between large and small males. We tested this with a PCA analysis (Figure S1c). PC1 explained 71.5% of the variation in the data. For PC1, we observed separate clustering based on male size, indicating that the composition of the seminal fluid proteome produced is different between large and small males when the data is not normalised (L $\text{ratio}^2_2 = 4.497$, $p = 0.033$). PC2 explained 9.3% of the variation in the data but this was not associated with the size of the male (L $\text{ratio}^2_2 = 0.719$, $p = 0.396$). Together, these results suggest that before the normalisation process, large males had significantly higher abundances of Sfps than small males and this difference was larger compared to the normalised data, but again these differences were not driven by any major changes in individual proteins.

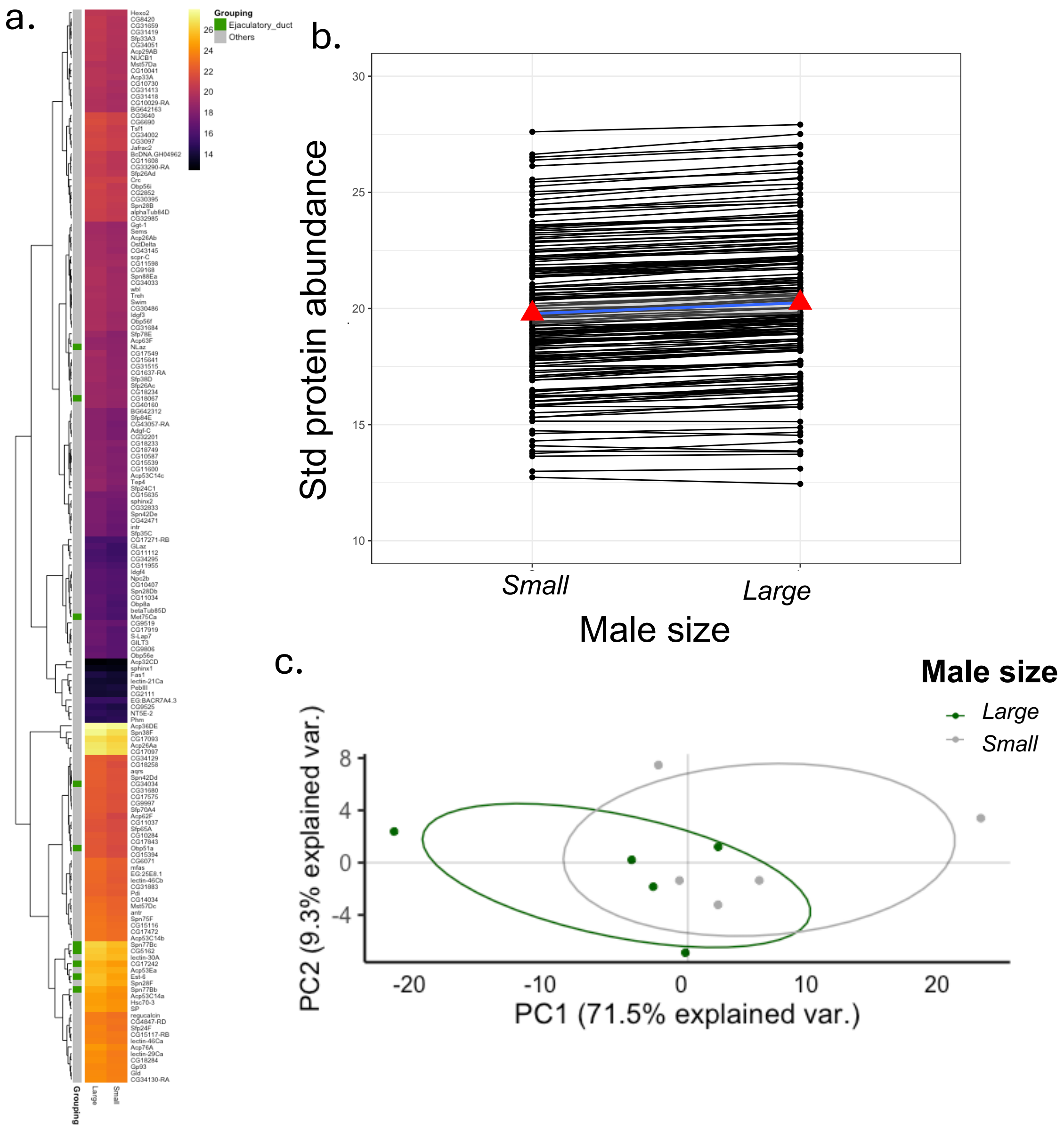


**Figure S1.** Seminal fluid protein production increases in large males compared to small males when analyses is repeated with unnormalized data. (A) Heatmap showing the abundance of 167 Sfps identified in accessory gland and ejaculatory duct samples of small and large males. Each cell gives the across-replicate mean for that Sfp in a given treatment (n=5 replicates per treatment). None of the Sfps changed in abundance as a response to male size after multiple test correction. Row annotation provide information relating to whether the Sfp is produced in the accessory glands or ejaculatory duct (green annotation). Pearson correlation was used as the distance metric for the hierarchical clustering. (B) Line plots showing the change in standardised Sfp abundance with male size. The mean standardised abundance of Sfps in small and large males are depicted with red triangles and joined by a blue line. Large males have a statistically significant increase in Sfp abundances compared to small males. (C) Principal component analyses of the seminal fluid proteome in male reproductive tissues. The composition of the seminal fluid proteome is different for small and large males. Red points represent small male virgin samples (5 replicates) and blue points represent large male virgin samples (5 replicates). Ellipses denote 80% normal probability.

The heatmap of transferred Sfps revealed that there was clustering among the Sfps in their response to male and female size (Figure S2a). We didn't analyse individual clusters here as they would be different group of Sfps compared to the clusters found from the normalised data but wanted to check if the general patterns were similar. Overall, small males transferred significantly higher abundances of Sfps than large males and more so to large females than small females (male size and female size interaction, L $ratio^2_2 = 9.781$, $p = 0.001$, Figure S2b). This agrees with the results from our main analyses and suggests that the results we find using the normalised data is not an artefact of the normalisation process.

Next, we looked at the composition of the seminal proteome transferred and again found it to be dependent on male size (Figure S2c). PC1 explained 37% of the variation in the data. For PC1, we observed separate clustering based on male size, indicating that the composition of the seminal fluid proteome transferred is different between large and small males (L $ratio^2_2 = 6.608$, $p = 0.01$). However, the composition of the seminal proteome transferred did not respond to female size or the interaction between male and female size (female size, L $ratio^2_2 = 0.561$, $p = 0.453$; interaction, L $ratio^2_2 = 0.386$, $p = 0.534$). PC2 explained 21.7% of the variation in the data but these were not associated with the size of the male, size of the female or their interaction (male size, L $ratio^2_2 = 2.107$, $p = 0.146$; female size, L $ratio^2_2 = 0.102$, $p = 0.748$; interaction, L $ratio^2_2 = 0.217$, $p = 0.64$).

Finally, we repeated the analyses to identify the key Sfps underpinning compositional effects. After correcting for multiple tests, we found that small males transferred 14 Sfps in significantly higher abundances compared to large males. Eight of these are among the 10 Sfps identified with the normalised data (Acp33A, CG10041, CG11955, CG17843, CG4847-RD, Obp56f, Phm, Sfp26Ad) and are transferred in significantly higher abundances by small males. Five other Sfps (Acp53Ea, CG31413, CG34129, PebIII, Sfp26Ac) are similarly transferred in higher abundances by small males but are only found in the analyses of the unnormalized data. Finally, Adgf-C was transferred in higher abundances by large males. Together, the results from the raw data suggest that small males transfer many Sfps in relatively higher abundances than large males and this result is not driven by the normalisation process itself as it is still evident without it.

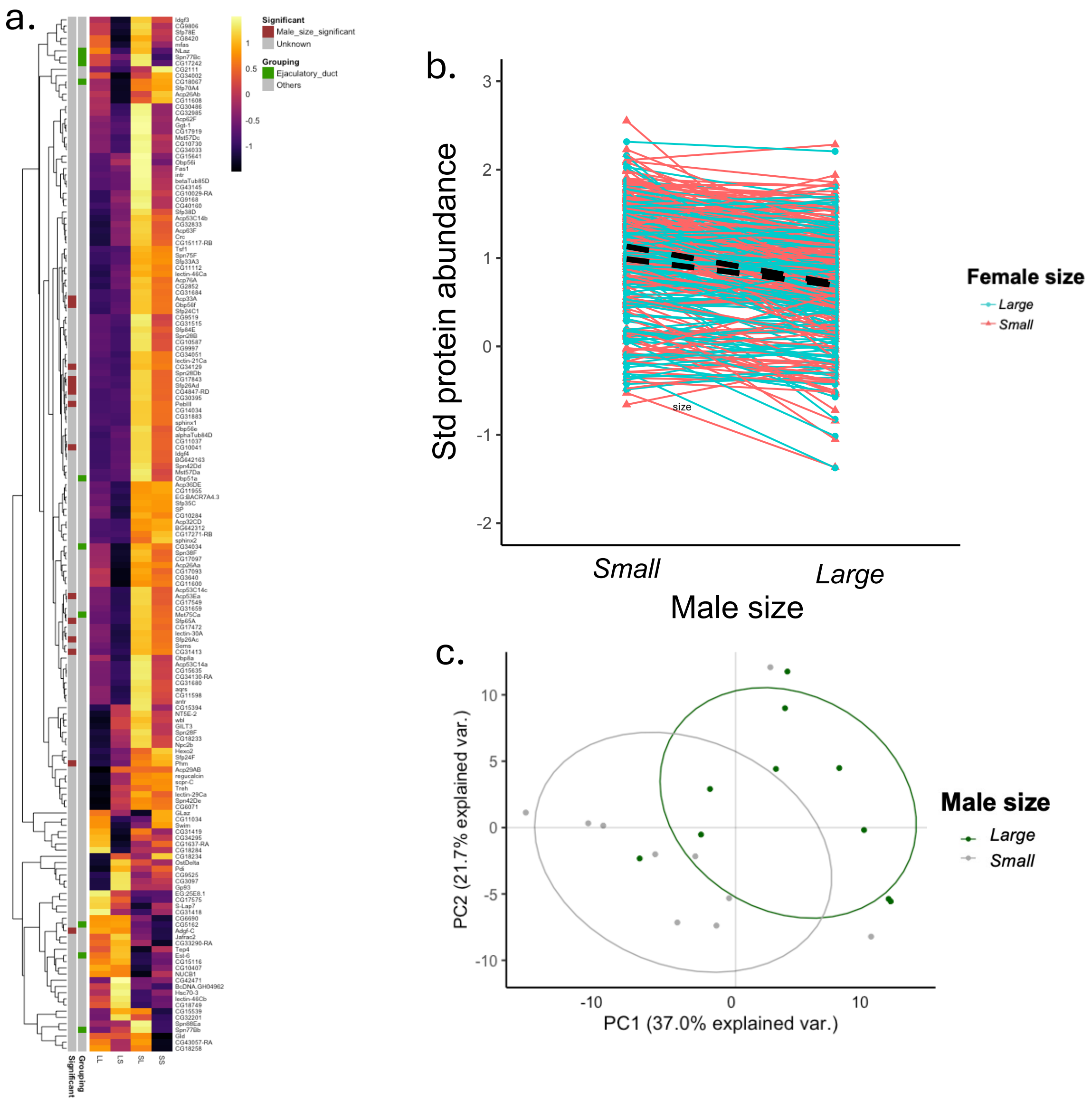


**Figure S2.** Seminal fluid protein transfer responds to male size when analyses are repeated with unnormalized data. (A) Heatmap showing the abundance of 167 Sfps transferred to females during mating (n=5 replicates per treatment). Large males were mated to large females (LL) or small females (LS) and small males were mated to large females (SL) or small females (SS). Each cell gives the across-replicate mean for that Sfp in a given treatment combination. 14 Sfps (Acp33A, CG10041, CG11955, CG17843, CG4847-RD, Obp56f, Phm, Sfp26Ad, Acp53Ea, CG31413, CG34129, PebIII, Sfp26Ac, Adgf-C) were transferred in significantly higher abundances by small males compared to large males. Row annotation provide information relating to whether the Sfp is produced in the accessory glands or ejaculatory duct (green annotation). Pearson correlation was used as the distance metric for the hierarchical clustering. (B) Line plots showing the standardised abundance of Sfps transferred as a response to male and female size. The average change in Sfp abundance for large and small females as a response to male size are depicted with dashed black lines marked as large F and small F. Small males transferred a higher abundance of these Sfps to large females. (C) Principal component analyses of the seminal fluid proteome transferred to females during mating. The composition of the seminal fluid proteome transferred is different for small and large males. Red points represent small male samples (5 replicates mated to small females and 5 replicates mated to large females – 10 in total) and blue points represent large male samples (5 replicates mated to small females and 5 replicates mated to large females – 10 in total). Ellipses denote 80% normal probability.

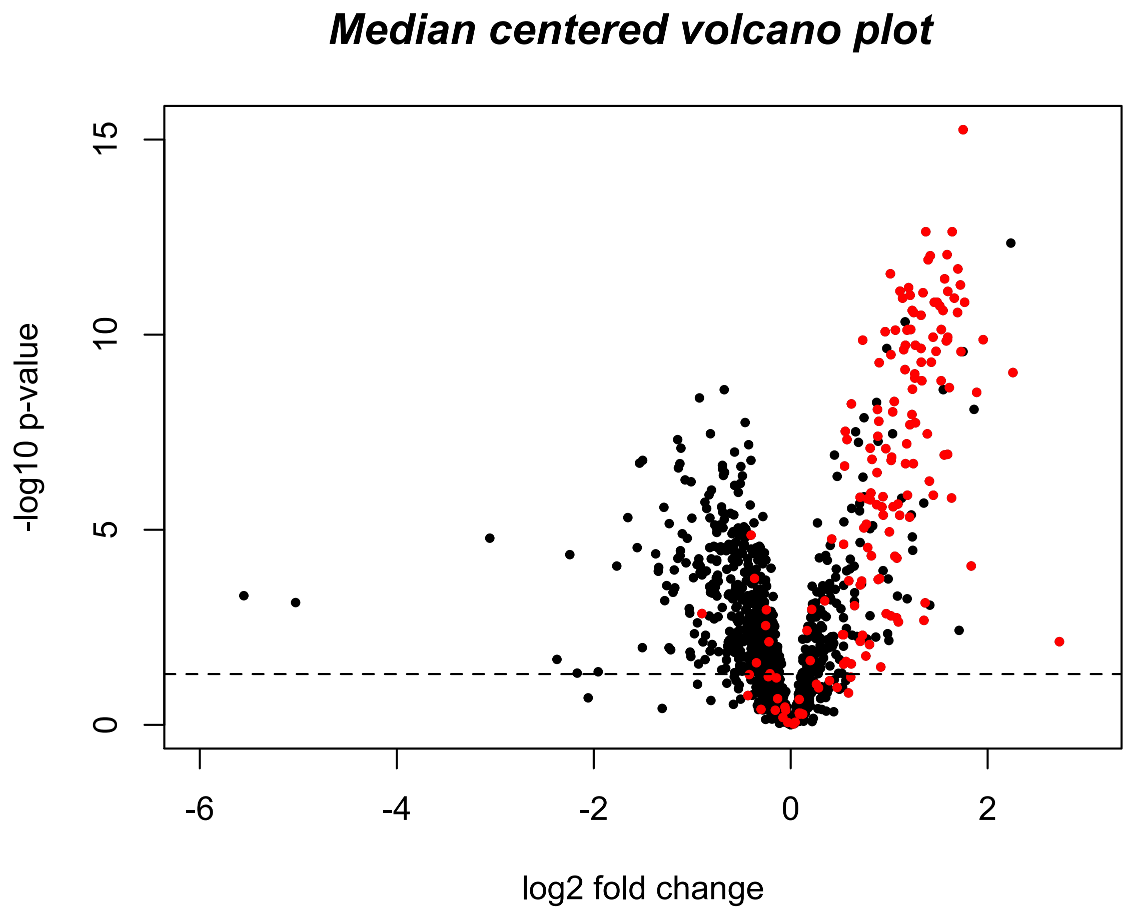


**Figure S3.** Volcano plot displaying all proteins detected that were identified by at least two unique peptides. The log2 fold change [log2 (virgin) - log2 (newly mated)] is shown on the *x* axis and significance is displayed on the *y* axis as the negative logarithm (log10 scale) of the fdr corrected *p* value. Known Sfps are coloured in red. The significance cutoff ($p < 0.05$) is highlighted with a dashed line.

**Supplementary Data S1.** The list of proteins detected in the study that were identified by at least two unique peptides and the data used to create the volcano plot displayed in Figure S3.

**Table S1.** The list of Sfps detected in this study. Each Sfps response to male size before and after FDR correction are included (*male size pval* and *male size qval* respectively).

| *Protein* | *male size pval* | *male size qval* |
|---|---|---|
| Acp26Aa | 0.4941 | 0.9942 |
| Acp26Ab | 0.7980 | 0.9942 |
| Acp29AB | 0.6715 | 0.9942 |
| Acp32CD | 0.4023 | 0.9942 |
| Acp33A | 0.6630 | 0.9942 |
| Acp36DE | 0.7800 | 0.9942 |
| Acp53C14a | 0.3670 | 0.9942 |
| Acp53C14b | 0.8106 | 0.9942 |
| Acp53C14c | 0.8291 | 0.9942 |
| Acp53Ea | 0.7327 | 0.9942 |
| Acp62F | 0.1460 | 0.9942 |
| Acp63F | 0.8460 | 0.9942 |
| Acp76A | 0.1645 | 0.9942 |
| Adgf-C | 0.2649 | 0.9942 |
| alphaTub84D | 0.9670 | 0.9942 |
| antr | 0.1867 | 0.9942 |
| aqrs | 0.3887 | 0.9942 |
| BcDNA.GH04962 | 0.1490 | 0.9942 |
| betaTub85D | 0.3617 | 0.9942 |
| BG642163 | 0.9531 | 0.9942 |
| BG642312 | 0.9838 | 0.9942 |
| CG10029-RA | 0.7035 | 0.9942 |
| CG10041 | 0.9942 | 0.9942 |
| CG10284 | 0.4814 | 0.9942 |
| CG10407 | 0.5588 | 0.9942 |
| CG10587 | 0.9455 | 0.9942 |
| CG10730 | 0.3001 | 0.9942 |
| CG11034 | 0.5154 | 0.9942 |
| CG11037 | 0.9795 | 0.9942 |
| CG11112 | 0.9264 | 0.9942 |
| CG11598 | 0.6325 | 0.9942 |
| CG11600 | 0.5583 | 0.9942 |
| CG11608 | 0.3960 | 0.9942 |
| CG11955 | 0.8483 | 0.9942 |
| CG14034 | 0.9665 | 0.9942 |
| CG15116 | 0.9939 | 0.9942 |
| CG15117-RB | 0.5303 | 0.9942 |
| CG15394 | 0.4828 | 0.9942 |
| CG15539 | 0.8505 | 0.9942 |
| CG15635 | 0.4550 | 0.9942 |
| CG15641 | 0.2405 | 0.9942 |
| CG1637-RA | 0.6746 | 0.9942 |
| CG17093 | 0.6709 | 0.9942 |
| CG17097 | 0.6431 | 0.9942 |
| CG17242 | 0.0470 | 0.9942 |

| | | |
|---|---|---|
| CG17271-RB | 0.3538 | 0.9942 |
| CG17472 | 0.6498 | 0.9942 |
| CG17549 | 0.0440 | 0.9942 |
| CG17575 | 0.5206 | 0.9942 |
| CG17843 | 0.9923 | 0.9942 |
| CG17919 | 0.3305 | 0.9942 |
| CG18067 | 0.8260 | 0.9942 |
| CG18233 | 0.2902 | 0.9942 |
| CG18234 | 0.2000 | 0.9942 |
| CG18258 | 0.1603 | 0.9942 |
| CG18284 | 0.2290 | 0.9942 |
| CG18749 | 0.5352 | 0.9942 |
| CG2111 | 0.2711 | 0.9942 |
| CG2852 | 0.9008 | 0.9942 |
| CG30395 | 0.7761 | 0.9942 |
| CG30486 | 0.7617 | 0.9942 |
| CG3097 | 0.9475 | 0.9942 |
| CG31413 | 0.9802 | 0.9942 |
| CG31418 | 0.4694 | 0.9942 |
| CG31419 | 0.5893 | 0.9942 |
| CG31515 | 0.2423 | 0.9942 |
| CG31659 | 0.1613 | 0.9942 |
| CG31680 | 0.7824 | 0.9942 |
| CG31684 | 0.5273 | 0.9942 |
| CG31883 | 0.7846 | 0.9942 |
| CG32201 | 0.0141 | 0.9942 |
| CG32833 | 0.7695 | 0.9942 |
| CG32985 | 0.4464 | 0.9942 |
| CG33290-RA | 0.7036 | 0.9942 |
| CG34002 | 0.7211 | 0.9942 |
| CG34033 | 0.3730 | 0.9942 |
| CG34034 | 0.7154 | 0.9942 |
| CG34051 | 0.9386 | 0.9942 |
| CG34129 | 0.3371 | 0.9942 |
| CG34130-RA | 0.9322 | 0.9942 |
| CG34295 | 0.8269 | 0.9942 |
| CG3640 | 0.8631 | 0.9942 |
| CG40160 | 0.7136 | 0.9942 |
| CG42471 | 0.5071 | 0.9942 |
| CG43057-RA | 0.1419 | 0.9942 |
| CG43145 | 0.5856 | 0.9942 |
| CG4847-RD | 0.7019 | 0.9942 |
| CG5162 | 0.0726 | 0.9942 |
| CG6071 | 0.4895 | 0.9942 |
| CG6690 | 0.5898 | 0.9942 |
| CG8420 | 0.4102 | 0.9942 |
| CG9168 | 0.1748 | 0.9942 |
| CG9519 | 0.5653 | 0.9942 |
| CG9525 | 0.9186 | 0.9942 |

| | | |
|---|---|---|
| CG9806 | 0.6985 | 0.9942 |
| CG9997 | 0.4886 | 0.9942 |
| Crc | 0.2098 | 0.9942 |
| EG:25E8.1 | 0.0315 | 0.9942 |
| EG:BACR7A4.3 | 0.2437 | 0.9942 |
| Est-6 | 0.1470 | 0.9942 |
| Fas1 | 0.8915 | 0.9942 |
| Ggt-1 | 0.8770 | 0.9942 |
| GILT3 | 0.0141 | 0.9942 |
| GLaz | 0.1417 | 0.9942 |
| Gld | 0.9055 | 0.9942 |
| Gp93 | 0.4841 | 0.9942 |
| Hexo2 | 0.2046 | 0.9942 |
| Hsc70-3 | 0.7081 | 0.9942 |
| Idgf3 | 0.2766 | 0.9942 |
| Idgf4 | 0.8887 | 0.9942 |
| intr | 0.1157 | 0.9942 |
| Jafrac2 | 0.6986 | 0.9942 |
| lectin-21Ca | 0.3108 | 0.9942 |
| lectin-29Ca | 0.1777 | 0.9942 |
| lectin-30A | 0.8322 | 0.9942 |
| lectin-46Ca | 0.7437 | 0.9942 |
| lectin-46Cb | 0.4475 | 0.9942 |
| Met75Ca | 0.5363 | 0.9942 |
| mfas | 0.2238 | 0.9942 |
| Mst57Da | 0.7438 | 0.9942 |
| Mst57Dc | 0.5256 | 0.9942 |
| NLaz | 0.6428 | 0.9942 |
| Npc2b | 0.9053 | 0.9942 |
| NT5E-2 | 0.5573 | 0.9942 |
| NUCB1 | 0.3538 | 0.9942 |
| Obp51a | 0.4658 | 0.9942 |
| Obp56e | 0.8791 | 0.9942 |
| Obp56f | 0.6609 | 0.9942 |
| Obp56i | 0.2268 | 0.9942 |
| Obp8a | 0.1419 | 0.9942 |
| OstDelta | 0.9695 | 0.9942 |
| Pdi | 0.6417 | 0.9942 |
| PebIII | 0.1171 | 0.9942 |
| Phm | 0.1987 | 0.9942 |
| regucalcin | 0.7169 | 0.9942 |
| S-Lap7 | 0.7035 | 0.9942 |
| scpr-C | 0.6364 | 0.9942 |
| Sems | 0.7806 | 0.9942 |
| Sfp24C1 | 0.2851 | 0.9942 |
| Sfp24F | 0.3792 | 0.9942 |
| Sfp26Ac | 0.9925 | 0.9942 |
| Sfp26Ad | 0.8987 | 0.9942 |
| Sfp33A3 | 0.5993 | 0.9942 |

| | | |
|---|---|---|
| Sfp35C | 0.6940 | 0.9942 |
| Sfp38D | 0.4088 | 0.9942 |
| Sfp65A | 0.7847 | 0.9942 |
| Sfp70A4 | 0.5359 | 0.9942 |
| Sfp78E | 0.3604 | 0.9942 |
| Sfp84E | 0.8541 | 0.9942 |
| SP | 0.9100 | 0.9942 |
| sphinx1 | 0.6934 | 0.9942 |
| sphinx2 | 0.9567 | 0.9942 |
| Spn28B | 0.6181 | 0.9942 |
| Spn28Db | 0.2641 | 0.9942 |
| Spn28F | 0.2019 | 0.9942 |
| Spn38F | 0.0886 | 0.9942 |
| Spn42Dd | 0.4930 | 0.9942 |
| Spn42De | 0.1221 | 0.9942 |
| Spn75F | 0.5323 | 0.9942 |
| Spn77Bb | 0.1836 | 0.9942 |
| Spn77Bc | 0.1176 | 0.9942 |
| Spn88Ea | 0.3433 | 0.9942 |
| Swim | 0.9051 | 0.9942 |
| Tep4 | 0.2780 | 0.9942 |
| Treh | 0.6239 | 0.9942 |
| Tsf1 | 0.1190 | 0.9942 |
| wbl | 0.7343 | 0.9942 |

**Table S2.** The list of Sfps detected in this study. The relative abundance of Sfps transferred as a response to male size, female size and their interaction before and after FDR correction are included (*male size pval*, *male size qval, female size pval*, f*emale size qval, male x female pval*, *male x female qval* respectively).

| *Protein* | *male size pval* | *male size qval* | *female size pval* | *female size qval* | *male x female pval* | *male x female qval* |
|---|---|---|---|---|---|---|
| Acp26Aa | 0.7059 | 0.7965 | 0.7589 | 0.9284 | 0.8427 | 0.9955 |
| Acp26Ab | 0.9357 | 0.9572 | 0.8315 | 0.9380 | 0.7192 | 0.9955 |
| Acp29AB | 0.2229 | 0.3799 | 0.8537 | 0.9380 | 0.8563 | 0.9955 |
| Acp32CD | 0.0140 | 0.0709 | 0.8363 | 0.9380 | 0.6790 | 0.9955 |
| Acp33A | 0.0011 | 0.0361 | 0.3400 | 0.8272 | 0.8487 | 0.9955 |
| Acp36DE | 0.0530 | 0.1404 | 0.9815 | 0.9859 | 0.8557 | 0.9955 |
| Acp53C14a | 0.0790 | 0.1816 | 0.1334 | 0.8272 | 0.7291 | 0.9955 |
| Acp53C14b | 0.0061 | 0.0601 | 0.4892 | 0.8272 | 0.7472 | 0.9955 |
| Acp53C14c | 0.0725 | 0.1781 | 0.1588 | 0.8272 | 0.6728 | 0.9955 |
| Acp53Ea | 0.0076 | 0.0613 | 0.1152 | 0.8272 | 0.6228 | 0.9955 |
| Acp62F | 0.6851 | 0.7875 | 0.3343 | 0.8272 | 0.9955 | 0.9955 |
| Acp63F | 0.0120 | 0.0703 | 0.4840 | 0.8272 | 0.6005 | 0.9955 |
| Acp76A | 0.0171 | 0.0818 | 0.6095 | 0.8355 | 0.5587 | 0.9955 |
| Adgf-C | 0.0077 | 0.0613 | 0.5318 | 0.8285 | 0.8360 | 0.9955 |
| alphaTub84D | 0.0526 | 0.1404 | 0.2907 | 0.8272 | 0.8963 | 0.9955 |
| antr | 0.1542 | 0.2893 | 0.1289 | 0.8272 | 0.5043 | 0.9955 |
| aqrs | 0.0767 | 0.1816 | 0.0644 | 0.8272 | 0.7005 | 0.9955 |
| BcDNA.GH04962 | 0.0901 | 0.1958 | 0.7761 | 0.9284 | 0.9872 | 0.9955 |
| betaTub85D | 0.6885 | 0.7875 | 0.4138 | 0.8272 | 0.7386 | 0.9955 |
| BG642163 | 0.0090 | 0.0654 | 0.2857 | 0.8272 | 0.8242 | 0.9955 |
| BG642312 | 0.0223 | 0.0925 | 0.9256 | 0.9573 | 0.8108 | 0.9955 |
| CG10029-RA | 0.0203 | 0.0894 | 0.1676 | 0.8272 | 0.6307 | 0.9955 |
| CG10041 | 0.0008 | 0.0361 | 0.1457 | 0.8272 | 0.2752 | 0.9955 |
| CG10284 | 0.1402 | 0.2723 | 0.5358 | 0.8285 | 0.7687 | 0.9955 |
| CG10407 | 0.0448 | 0.1247 | 0.8304 | 0.9380 | 0.5915 | 0.9955 |
| CG10587 | 0.0181 | 0.0839 | 0.1229 | 0.8272 | 0.9880 | 0.9955 |
| CG10730 | 0.4198 | 0.5711 | 0.2318 | 0.8272 | 0.7827 | 0.9955 |
| CG11034 | 0.6418 | 0.7602 | 0.7745 | 0.9284 | 0.8384 | 0.9955 |
| CG11037 | 0.0298 | 0.1036 | 0.4060 | 0.8272 | 0.8083 | 0.9955 |
| CG11112 | 0.0419 | 0.1218 | 0.6241 | 0.8473 | 0.9210 | 0.9955 |
| CG11598 | 0.2583 | 0.4229 | 0.3627 | 0.8272 | 0.9234 | 0.9955 |
| CG11600 | 0.9287 | 0.9572 | 0.9290 | 0.9573 | 0.9604 | 0.9955 |
| CG11608 | 0.3745 | 0.5366 | 0.8524 | 0.9380 | 0.4068 | 0.9955 |
| CG11955 | 0.0011 | 0.0361 | 0.9344 | 0.9573 | 0.8336 | 0.9955 |
| CG14034 | 0.0073 | 0.0613 | 0.3047 | 0.8272 | 0.9563 | 0.9955 |
| CG15116 | 0.1690 | 0.3102 | 0.5935 | 0.8355 | 0.4465 | 0.9955 |
| CG15117-RB | 0.0123 | 0.0703 | 0.3814 | 0.8272 | 0.7543 | 0.9955 |
| CG15394 | 0.0351 | 0.1107 | 0.0977 | 0.8272 | 0.4074 | 0.9955 |

| | | | | | | |
|---|---|---|---|---|---|---|
| CG15539 | 0.5251 | 0.6645 | 0.5357 | 0.8285 | 0.9233 | 0.9955 |
| CG15635 | 0.0126 | 0.0703 | 0.0544 | 0.8272 | 0.7198 | 0.9955 |
| CG15641 | 0.2468 | 0.4080 | 0.1377 | 0.8272 | 0.2417 | 0.9955 |
| CG1637-RA | 0.1910 | 0.3392 | 0.6900 | 0.9002 | 0.3068 | 0.9955 |
| CG17093 | 0.2059 | 0.3545 | 0.1844 | 0.8272 | 0.5263 | 0.9955 |
| CG17097 | 0.2004 | 0.3486 | 0.4766 | 0.8272 | 0.9585 | 0.9955 |
| CG17242 | 0.3209 | 0.4829 | 0.7496 | 0.9284 | 0.7998 | 0.9955 |
| CG17271-RB | 0.0227 | 0.0925 | 0.9859 | 0.9859 | 0.5426 | 0.9955 |
| CG17472 | 0.1315 | 0.2584 | 0.3706 | 0.8272 | 0.9520 | 0.9955 |
| CG17549 | 0.6847 | 0.7875 | 0.6104 | 0.8355 | 0.9620 | 0.9955 |
| CG17575 | 0.1187 | 0.2389 | 0.5603 | 0.8292 | 0.4648 | 0.9955 |
| CG17843 | 0.0014 | 0.0391 | 0.1476 | 0.8272 | 0.7534 | 0.9955 |
| CG17919 | 0.3858 | 0.5460 | 0.2669 | 0.8272 | 0.9532 | 0.9955 |
| CG18067 | 0.0391 | 0.1166 | 0.7158 | 0.9237 | 0.3572 | 0.9955 |
| CG18233 | 0.2363 | 0.3946 | 0.4801 | 0.8272 | 0.7855 | 0.9955 |
| CG18234 | 0.3976 | 0.5576 | 0.4999 | 0.8272 | 0.3827 | 0.9955 |
| CG18258 | 0.2757 | 0.4384 | 0.2984 | 0.8272 | 0.7475 | 0.9955 |
| CG18284 | 0.3023 | 0.4674 | 0.6427 | 0.8655 | 0.6168 | 0.9955 |
| CG18749 | 0.0100 | 0.0676 | 0.5759 | 0.8292 | 0.9142 | 0.9955 |
| CG2111 | 0.2294 | 0.3869 | 0.4518 | 0.8272 | 0.3333 | 0.9955 |
| CG2852 | 0.0346 | 0.1107 | 0.5555 | 0.8292 | 0.9895 | 0.9955 |
| CG30395 | 0.0089 | 0.0654 | 0.3238 | 0.8272 | 0.7872 | 0.9955 |
| CG30486 | 0.5896 | 0.7135 | 0.1374 | 0.8272 | 0.7068 | 0.9955 |
| CG3097 | 0.5302 | 0.6645 | 0.5312 | 0.8285 | 0.6427 | 0.9955 |
| CG31413 | 0.0351 | 0.1107 | 0.4177 | 0.8272 | 0.8609 | 0.9955 |
| CG31418 | 0.1125 | 0.2348 | 0.9430 | 0.9603 | 0.6154 | 0.9955 |
| CG31419 | 0.4007 | 0.5576 | 0.5717 | 0.8292 | 0.8883 | 0.9955 |
| CG31515 | 0.0129 | 0.0703 | 0.0857 | 0.8272 | 0.5729 | 0.9955 |
| CG31659 | 0.0614 | 0.1553 | 0.2020 | 0.8272 | 0.6993 | 0.9955 |
| CG31680 | 0.0291 | 0.1033 | 0.0921 | 0.8272 | 0.7643 | 0.9955 |
| CG31684 | 0.0076 | 0.0613 | 0.5082 | 0.8272 | 0.6536 | 0.9955 |
| CG31883 | 0.0775 | 0.1816 | 0.4483 | 0.8272 | 0.9618 | 0.9955 |
| CG32201 | 0.3664 | 0.5321 | 0.4498 | 0.8272 | 0.8374 | 0.9955 |
| CG32833 | 0.0794 | 0.1816 | 0.5661 | 0.8292 | 0.7770 | 0.9955 |
| CG32985 | 0.7267 | 0.8090 | 0.0591 | 0.8272 | 0.9428 | 0.9955 |
| CG33290-RA | 0.4178 | 0.5711 | 0.9037 | 0.9543 | 0.9300 | 0.9955 |
| CG34002 | 0.9103 | 0.9501 | 0.7364 | 0.9284 | 0.3449 | 0.9955 |
| CG34033 | 0.1437 | 0.2758 | 0.1187 | 0.8272 | 0.6477 | 0.9955 |
| CG34034 | 0.4267 | 0.5711 | 0.5532 | 0.8292 | 0.7541 | 0.9955 |
| CG34051 | 0.0133 | 0.0703 | 0.2833 | 0.8272 | 0.8640 | 0.9955 |
| CG34129 | 0.0052 | 0.0601 | 0.4631 | 0.8272 | 0.8261 | 0.9955 |
| CG34130-RA | 0.1174 | 0.2389 | 0.1453 | 0.8272 | 0.7228 | 0.9955 |
| CG34295 | 0.7715 | 0.8366 | 0.0614 | 0.8272 | 0.1090 | 0.9955 |
| CG3640 | 0.1057 | 0.2234 | 0.1734 | 0.8272 | 0.3589 | 0.9955 |
| CG40160 | 0.0290 | 0.1033 | 0.1888 | 0.8272 | 0.0717 | 0.9955 |
| CG42471 | 0.3280 | 0.4890 | 0.3022 | 0.8272 | 0.3650 | 0.9955 |

| | | | | | | |
|---|---|---|---|---|---|---|
| CG43057-RA | 0.2631 | 0.4266 | 0.0305 | 0.8272 | 0.7660 | 0.9955 |
| CG43145 | 0.0903 | 0.1958 | 0.2146 | 0.8272 | 0.4111 | 0.9955 |
| CG4847-RD | 0.0028 | 0.0471 | 0.3085 | 0.8272 | 0.7063 | 0.9955 |
| CG5162 | 0.0135 | 0.0703 | 0.7190 | 0.9237 | 0.9456 | 0.9955 |
| CG6071 | 0.3013 | 0.4674 | 0.9062 | 0.9543 | 0.8095 | 0.9955 |
| CG6690 | 0.5090 | 0.6539 | 0.8763 | 0.9473 | 0.9588 | 0.9955 |
| CG8420 | 0.9167 | 0.9509 | 0.1689 | 0.8272 | 0.6231 | 0.9955 |
| CG9168 | 0.5299 | 0.6645 | 0.5679 | 0.8292 | 0.6945 | 0.9955 |
| CG9519 | 0.3407 | 0.5034 | 0.4619 | 0.8272 | 0.4722 | 0.9955 |
| CG9525 | 0.6418 | 0.7602 | 0.0784 | 0.8272 | 0.3819 | 0.9955 |
| CG9806 | 0.4819 | 0.6287 | 0.1972 | 0.8272 | 0.9594 | 0.9955 |
| CG9997 | 0.6204 | 0.7454 | 0.4925 | 0.8272 | 0.9718 | 0.9955 |
| Crc | 0.0323 | 0.1078 | 0.5107 | 0.8272 | 0.8200 | 0.9955 |
| EG:25E8.1 | 0.0249 | 0.0942 | 0.4707 | 0.8272 | 0.4362 | 0.9955 |
| EG:BACR7A4.3 | 0.0122 | 0.0703 | 0.6814 | 0.8984 | 0.7029 | 0.9955 |
| Est-6 | 0.0964 | 0.2065 | 0.7659 | 0.9284 | 0.8529 | 0.9955 |
| Fas1 | 0.3473 | 0.5088 | 0.4386 | 0.8272 | 0.8167 | 0.9955 |
| Ggt-1 | 0.4547 | 0.5979 | 0.1634 | 0.8272 | 0.8838 | 0.9955 |
| GILT3 | 0.5493 | 0.6745 | 0.6074 | 0.8355 | 0.0195 | 0.9955 |
| GLaz | 0.6717 | 0.7875 | 0.9330 | 0.9573 | 0.5225 | 0.9955 |
| Gld | 0.3199 | 0.4829 | 0.1800 | 0.8272 | 0.6239 | 0.9955 |
| Gp93 | 0.1896 | 0.3392 | 0.4446 | 0.8272 | 0.4938 | 0.9955 |
| Hexo2 | 0.0773 | 0.1816 | 0.3056 | 0.8272 | 0.9353 | 0.9955 |
| Hsc70-3 | 0.0839 | 0.1869 | 0.6832 | 0.8984 | 0.7249 | 0.9955 |
| Idgf3 | 0.8583 | 0.9130 | 0.4127 | 0.8272 | 0.8999 | 0.9955 |
| Idgf4 | 0.0254 | 0.0942 | 0.4195 | 0.8272 | 0.6869 | 0.9955 |
| intr | 0.4277 | 0.5711 | 0.3844 | 0.8272 | 0.3632 | 0.9955 |
| Jafrac2 | 0.0423 | 0.1218 | 0.7796 | 0.9284 | 0.8141 | 0.9955 |
| lectin-21Ca | 0.0058 | 0.0601 | 0.2998 | 0.8272 | 0.8568 | 0.9955 |
| lectin-29Ca | 0.6934 | 0.7878 | 0.8107 | 0.9380 | 0.5852 | 0.9955 |
| lectin-30A | 0.0604 | 0.1551 | 0.2854 | 0.8272 | 0.7784 | 0.9955 |
| lectin-46Ca | 0.7592 | 0.8287 | 0.8831 | 0.9473 | 0.9066 | 0.9955 |
| lectin-46Cb | 0.2695 | 0.4328 | 0.8849 | 0.9473 | 0.9684 | 0.9955 |
| Met75Ca | 0.1563 | 0.2901 | 0.4204 | 0.8272 | 0.8928 | 0.9955 |
| mfas | 0.9400 | 0.9572 | 0.1431 | 0.8272 | 0.9636 | 0.9955 |
| Mst57Da | 0.0370 | 0.1140 | 0.2962 | 0.8272 | 0.5772 | 0.9955 |
| Mst57Dc | 0.1488 | 0.2824 | 0.0668 | 0.8272 | 0.5884 | 0.9955 |
| NLaz | 0.4925 | 0.6376 | 0.1469 | 0.8272 | 0.7286 | 0.9955 |
| Npc2b | 0.1975 | 0.3471 | 0.4448 | 0.8272 | 0.9667 | 0.9955 |
| NT5E-2 | 0.9897 | 0.9911 | 0.8360 | 0.9380 | 0.9521 | 0.9955 |
| NUCB1 | 0.0562 | 0.1465 | 0.4321 | 0.8272 | 0.6358 | 0.9955 |
| Obp51a | 0.0839 | 0.1869 | 0.3673 | 0.8272 | 0.8299 | 0.9955 |
| Obp56e | 0.0224 | 0.0925 | 0.3294 | 0.8272 | 0.7539 | 0.9955 |
| Obp56f | 0.0016 | 0.0393 | 0.3996 | 0.8272 | 0.8038 | 0.9955 |
| Obp56i | 0.7864 | 0.8472 | 0.4170 | 0.8272 | 0.3721 | 0.9955 |
| Obp8a | 0.5896 | 0.7135 | 0.2522 | 0.8272 | 0.8504 | 0.9955 |
| OstDelta | 0.5332 | 0.6645 | 0.7621 | 0.9284 | 0.7808 | 0.9955 |
| Pdi | 0.7348 | 0.8127 | 0.9501 | 0.9616 | 0.6741 | 0.9955 |

| | | | | | | |
|---|---|---|---|---|---|---|
| PebIII | 0.0050 | 0.0601 | 0.3333 | 0.8272 | 0.9663 | 0.9955 |
| Phm | 0.0001 | 0.0097 | 0.7839 | 0.9284 | 0.6119 | 0.9955 |
| regucalcin | 0.0008 | 0.0361 | 0.4328 | 0.8272 | 0.6995 | 0.9955 |
| S-Lap7 | 0.2820 | 0.4443 | 0.6754 | 0.8984 | 0.9947 | 0.9955 |
| scpr-C | 0.0441 | 0.1247 | 0.8473 | 0.9380 | 0.8449 | 0.9955 |
| Sems | 0.0244 | 0.0942 | 0.2402 | 0.8272 | 0.6844 | 0.9955 |
| Sfp24C1 | 0.0234 | 0.0932 | 0.4625 | 0.8272 | 0.9381 | 0.9955 |
| Sfp24F | 0.9911 | 0.9911 | 0.8013 | 0.9380 | 0.9447 | 0.9955 |
| Sfp26Ac | 0.0101 | 0.0676 | 0.1438 | 0.8272 | 0.6067 | 0.9955 |
| Sfp26Ad | 0.0019 | 0.0404 | 0.1702 | 0.8272 | 0.8003 | 0.9955 |
| Sfp33A3 | 0.0028 | 0.0471 | 0.5152 | 0.8272 | 0.7123 | 0.9955 |
| Sfp35C | 0.1139 | 0.2349 | 0.5051 | 0.8272 | 0.5453 | 0.9955 |
| Sfp38D | 0.0468 | 0.1282 | 0.4021 | 0.8272 | 0.6492 | 0.9955 |
| Sfp65A | 0.0060 | 0.0601 | 0.1953 | 0.8272 | 0.8675 | 0.9955 |
| Sfp70A4 | 0.7225 | 0.8090 | 0.8524 | 0.9380 | 0.7359 | 0.9955 |
| Sfp78E | 0.9763 | 0.9882 | 0.5981 | 0.8355 | 0.9021 | 0.9955 |
| Sfp84E | 0.1709 | 0.3102 | 0.4188 | 0.8272 | 0.9287 | 0.9955 |
| SP | 0.3090 | 0.4735 | 0.9086 | 0.9543 | 0.9180 | 0.9955 |
| sphinx1 | 0.1270 | 0.2525 | 0.5835 | 0.8329 | 0.9727 | 0.9955 |
| sphinx2 | 0.4247 | 0.5711 | 0.7448 | 0.9284 | 0.8221 | 0.9955 |
| Spn28B | 0.0148 | 0.0725 | 0.0367 | 0.8272 | 0.8375 | 0.9955 |
| Spn28Db | 0.0034 | 0.0513 | 0.1078 | 0.8272 | 0.9765 | 0.9955 |
| Spn28F | 0.0307 | 0.1045 | 0.1677 | 0.8272 | 0.8509 | 0.9955 |
| Spn38F | 0.8876 | 0.9323 | 0.7471 | 0.9284 | 0.9622 | 0.9955 |
| Spn42Dd | 0.0043 | 0.0601 | 0.1443 | 0.8272 | 0.4668 | 0.9955 |
| Spn42De | 0.7962 | 0.8523 | 0.4951 | 0.8272 | 0.4413 | 0.9955 |
| Spn75F | 0.0375 | 0.1140 | 0.4979 | 0.8272 | 0.7301 | 0.9955 |
| Spn77Bb | 0.8844 | 0.9323 | 0.1587 | 0.8272 | 0.2006 | 0.9955 |
| Spn77Bc | 0.7440 | 0.8175 | 0.2567 | 0.8272 | 0.8007 | 0.9955 |
| Spn88Ea | 0.5373 | 0.6646 | 0.4055 | 0.8272 | 0.7608 | 0.9955 |
| Swim | 0.6794 | 0.7875 | 0.8381 | 0.9380 | 0.1509 | 0.9955 |
| Tep4 | 0.0644 | 0.1606 | 0.2290 | 0.8272 | 0.8999 | 0.9955 |
| Treh | 0.3759 | 0.5366 | 0.8622 | 0.9411 | 0.7189 | 0.9955 |
| Tsf1 | 0.0200 | 0.0894 | 0.5578 | 0.8292 | 0.9359 | 0.9955 |
| wbl | 0.4309 | 0.5711 | 0.4778 | 0.8272 | 0.8702 | 0.9955 |
